# *In Silico* Epitope Prediction And VP1 Modelling For Foot-And-Mouth Serotype SAT 2 For Vaccine Design In East Africa

**DOI:** 10.1101/2021.09.13.460008

**Authors:** Jean Claude Udahemuka, Lunayo Accadius, George Obiero, Gabriel Aboge, Phiyani Lebea

**Affiliations:** Centre for Biotechnology and Bioinformatics, University of Nairobi, P.O. Box 30197, Nairobi, Kenya; Department of Veterinary Medicine, University of Rwanda, P.O. Box 57, Nyagatare, Rwanda; Department of Public Health, Pharmacology and Toxicology, University of Nairobi, P.O. Box 29053, Nairobi, Kenya; TokaBio (Pty), Ltd, Pretoria, South Africa

**Keywords:** FMDV, SAT2, Epitope prediction, Vaccine design, VP1 variability

## Abstract

Foot and Mouth Disease Virus has seven distinct, geographically localized, serotypes and a vaccination targeting one serotype does not confer immunity against another serotype. The use of inactivated vaccines is not safe and confers an immunity with a relatively shorter time. Using the VP1 sequences isolated in East Africa, we have predicted epitopes able to induce humoral and cell-mediated immunity in cattle. The Wu-Kabat variability index calculated in this study reflects the variable, including the known GH loop, and conserved regions, with the latter being good candidates for region-tailored vaccine design. Furthermore, we modelled the identified epitopes on a 3D model (PDB ID:5aca) to represent the epitopes structurally. This study can be used for *in vitro* and *in vivo* experiments.

## 1. Introduction

Foot-and-Mouth Disease (FMD) is an infectious disease caused by the Foot-and-Mouth Disease Virus (FMDV) of the family of picornaviridae (1). FMD is endemic in Africa and South-Eastern Asia with sporadic outbreaks in other countries (2–4). Seven distinct serotypes have been identified and geographically localized with seven pools (4) and SAT2 has been responsible for several outbreaks in Eastern Rwanda (5). Vaccination against one serotype does not confer immunity against other serotypes (6). Therefore, high potency multivalent vaccines have been developed (7) and a continuous study of molecular epidemiology at the regional and national level is of paramount importance to have regional-tailored vaccines. Poor understanding of circulating strains leads to limited vaccine matching and vaccine failure (8). The peptide-based vaccines paradigm has been proposed as an alternative to the currently used inactivated vaccines (9–11). This study is exploring the variability of FMDV SAT2 VP1 protein in EAC and *in silico* predicting highly potential epitopes to be targeted for epitope-based vaccines. Furthermore, we have complemented the prediction with a three-dimension (3D) presentation of the FMDV capsid using a close model available on the Protein Database Bank platform. The epitopes predicted in this study can be used to guide in choosing strains to include in vaccines or designing a peptide-based vaccine.

## 2. Materials and Methods

### 2.1. Selection of FMDV SAT2 VP1 sequences

In this study’s analysis, we analyzed the sequences from the field samples we collected and other sequences available online. For the latter, we searched online on three repositories namely; NCBI, EBI and DDBJ, using the following criteria, sequences identified as SAT2, sequences of VP1, sequence length between 212 and 216 amino acids, sequences of samples collected in the FMD pool IV, Great lakes region cluster countries (Burundi, Kenya, Rwanda, Tanzania and Uganda) together with DRC and samples collected after the year 2010. The latter criteria were not followed for samples collected in Burundi, DRC and Rwanda because of the scarcity of available sequences. The selected sequences following the above criteria are available online at 10.6084/m9.figshare.14979594.

### 2.2. VP1 sequence variability in Great Lakes cluster of FMDV pool IV

We used a Python 3 notebook to create a dictionary that can transcribe the bases of the raw field sequences into amino acids. The EBI ClustalW was used to run a multiple sequence alignment to obtain a consensus sequence. The VP1 consensus sequence was used to search for a three-dimensional (3D) model of the structure that is more likely to be found in the region of East Africa. The protein variability server (PVS) tool was used to calculate and represent a Wu-Kabat coefficient for variability following the formula below:

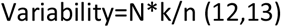

With N being the number of sequences in the alignment, k is the number of different amino acids at each position and n is the number of times for the most common amino acids per position. The Wu-Kabat coefficient was used to compare the variability of the predicted epitopes.

### 2.3. Epitope prediction and 3D modelling

The PVS was used to run a sequence alignment and analyze the variability of the selected sequences. To evaluate the positional variability of amino acids, we used the Wu-Kabat coefficient. SWISS-MODEL was used to select the fittest online available model and run its evaluation. The stereochemistry of the structure was evaluated using the Ramachandran plot to study the acceptability of the model.

We used the BepiPred-2.0 server to predict sequential B-cell epitopes with an epitope threshold set at 0.5. Likewise, the MHC-I binding predictions were made on 7/28/2021 using the IEDB analysis resource NetMHCpan (ver. 4.1) tool (14). For the protein digestion, we set the IEDB recommended 2020.09 (NetMHCpan EL 4.1) settings and digesting the protein in peptides ranging from 8 to 14-mers. The Bovine Leukocyte Antigen (BoLA) alleles available on the platform were all selected as ligands with default parameters. The peptides scoring more than 0.5 were considered for further analysis. The PyMol 2.5 (15) was used for the visualization of the selected model and to map the identified epitopes on the FMDV VP1 in 3-D presentation.

## 3. Results

### 3.1. VP1 sequence variability in Great Lakes cluster of FMDV pool IV

We used Python 3 to transcribe field sequence bases into amino acids and aligned field sequences with retrieved sequences. The VP1 sequences alignment using the CLC Main Work Bench 21.0.4 is presented in figure 1. The protein variability plot was obtained using the Protein Variability Server (PVS) by calculating the Wu-Kabat coefficient Figure 2. We identified at least two regions (residue 45-60 and residue 138-140) that are highly variable across the selected sequences and position 58 being the most variable.

**Figure 1:**
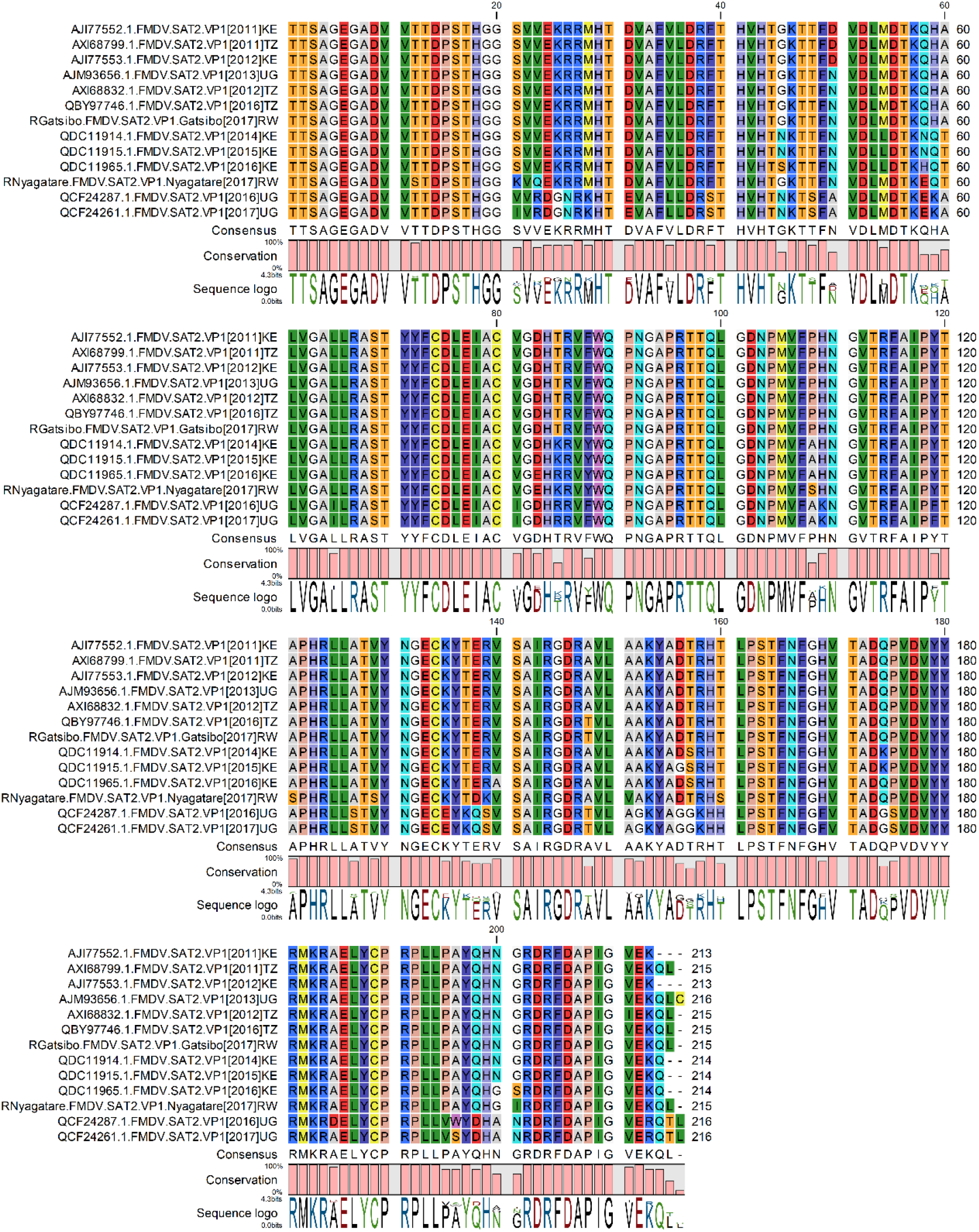
Similarity alignment of putative VP1 proteins of Foot and Mouth Disease serotype SAT 2 in East Africa

**Figure 2:**
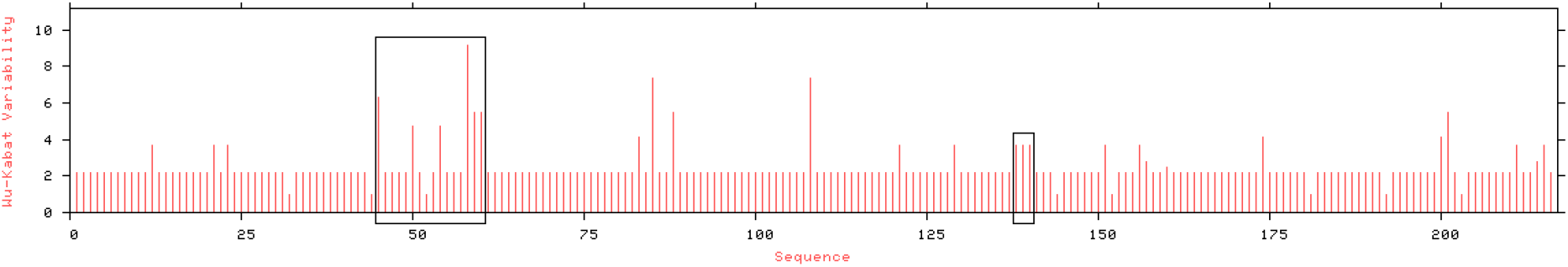
Wu-Kabat plot displaying the variability of VP1 proteins of FMDV SAT2 isolates from East Africa. The black rectangles highlight variable motifs with at least 3 variable positions. The plot was constructed using the Protein Variability Server (13).

The residue 58-60 and at positions 85, 108 were variable in a way that Nyagatare isolate is identical to isolates QDC11914, QDC11915 and QDC11965 isolated in cattle (*Bos taurus)* in Kenya at the livestock-wildlife interface between 214 and 2016. Likewise, at the same positions, the Gatsibo isolate was identical to AJI77552 (cattle, Kenya, 2011), AJI77553 (cattle, Kenya, 2012), AJM93656 (cattle, Uganda, 2013), AXI68799 (cattle, Tanzania, 2011), AXI68832 (cattle, Tanzania, 2012) and QBY97746 (cattle, Tanzania, 2016). The below phylogenetic representation (Figure 3) illustrates that at least FMD SAT2 viruses circulating in the region can be considered as two bigger branches.

**Figure 3:**
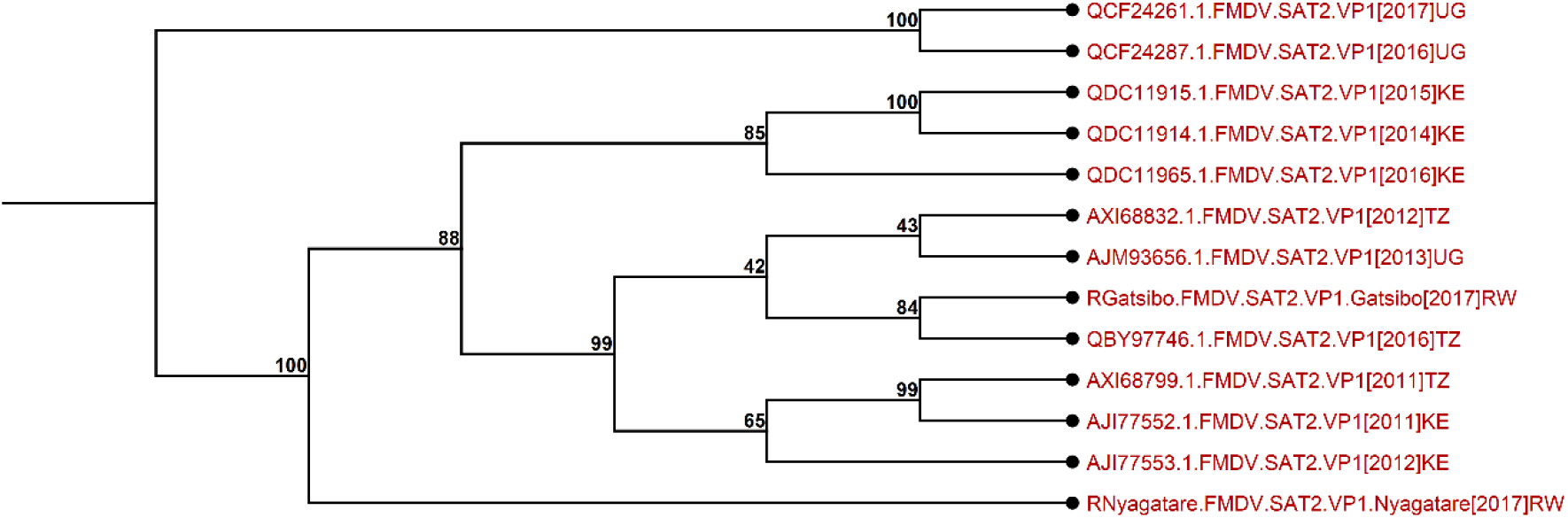
Phylogeny of FMD SAT2 isolates circulating in East Africa

### 3.2. Epitope prediction and 3D modelling

The multiple sequence alignment of the 11 FMD VP1 sequences was obtained using the European Bioinformatics Institute (EBI) online platform. Using the EBI’s EMBOSS program, the resulting consensus sequence (TTSAGEGADVVTTDPSTHGGSVVEKRRMHTDVAFVLDRFTHVHTGKTTFNVDLMDTKQHALVGALLRASTYYFCDLEIACVGDHTRVFWQPNGAPRTTQLGDNPMVFPHNGVTRFAIPYTAPHRLLATVYNGECKYTERVSAIRGDRAVLAAKYADTRHTLPSTFNFGHVTADQPVDVYYRMKRAELYCPRPLLPAYQHNGRDRFDAPIGVEKQLC) was obtained. We searched for a model close to the consensus sequence with available structures in the protein data bank using the SWISS-MODEL, the model with PDB id: 5ACA was selected. The modelled structure had a global QMEANDisCo of (0.77± 0.06) with a molprobity score of 1.50 and a Ramachandran favoured score of 91.35% A Ramachandran plot (Figure 4) showed that the structure can be accepted and considered for 3D structure presentation.

**Figure 4:**
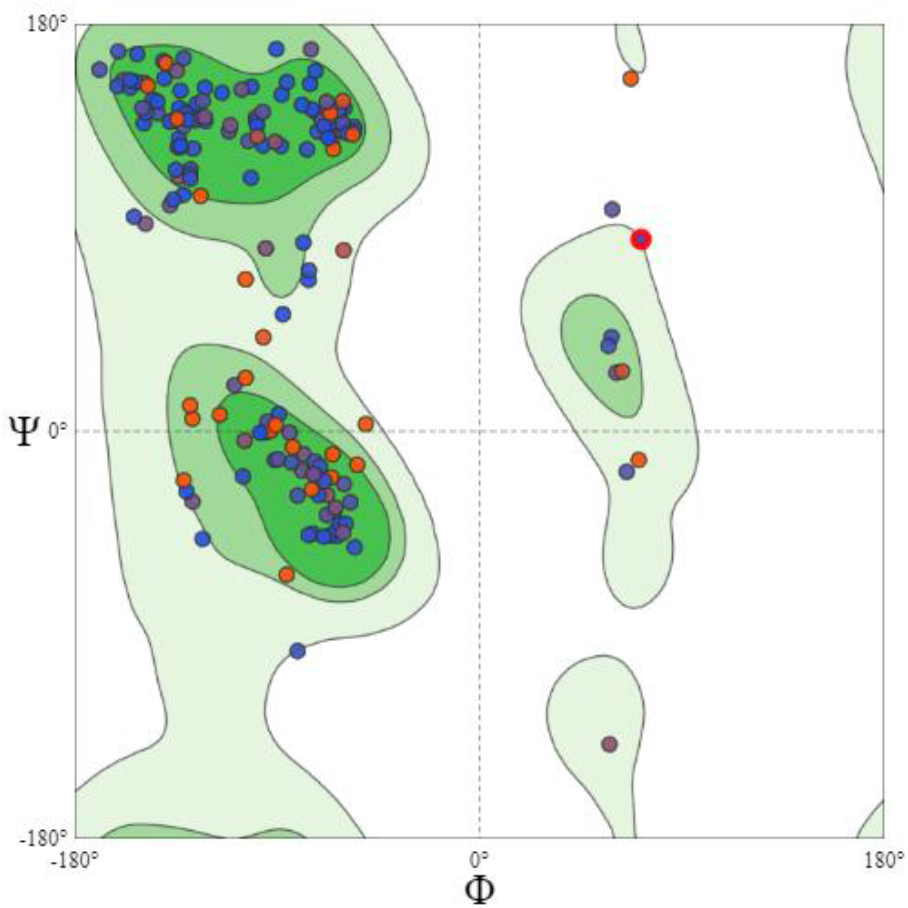
A Ramachandran plot for the selected 3D structure model showing the position of different amino acids

#### 3.2.1. B-cell epitopes

The mapping of epitopes in Figure 5 was obtained with the consensus sequence to predict the sequential B-cell epitopes DTU online platform (BepiPred-2.0) (16) and Table 1 describes the predicted peptides with their positions.

**Table 1:**
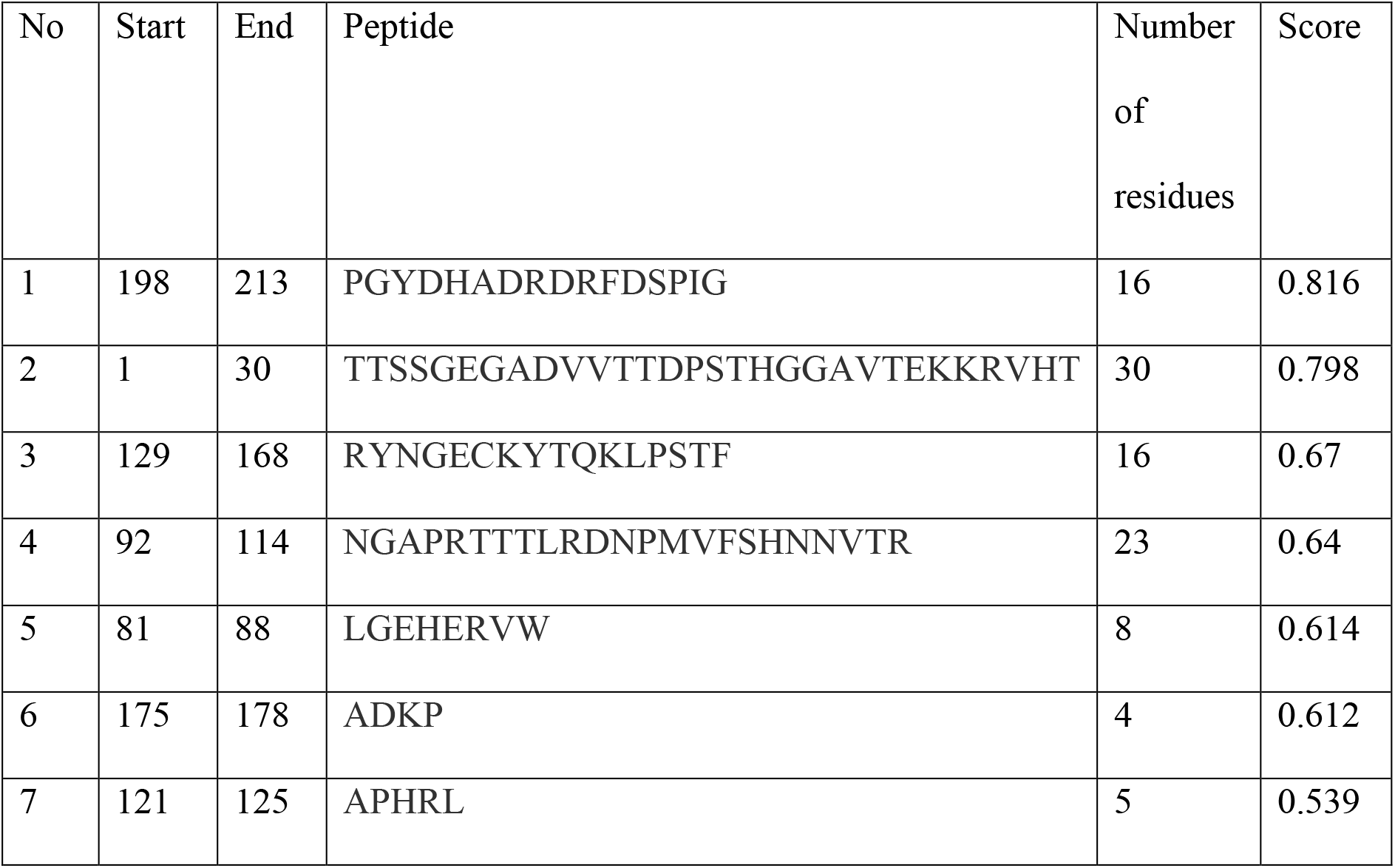
Predicted linear epitopes

**Figure 5:**
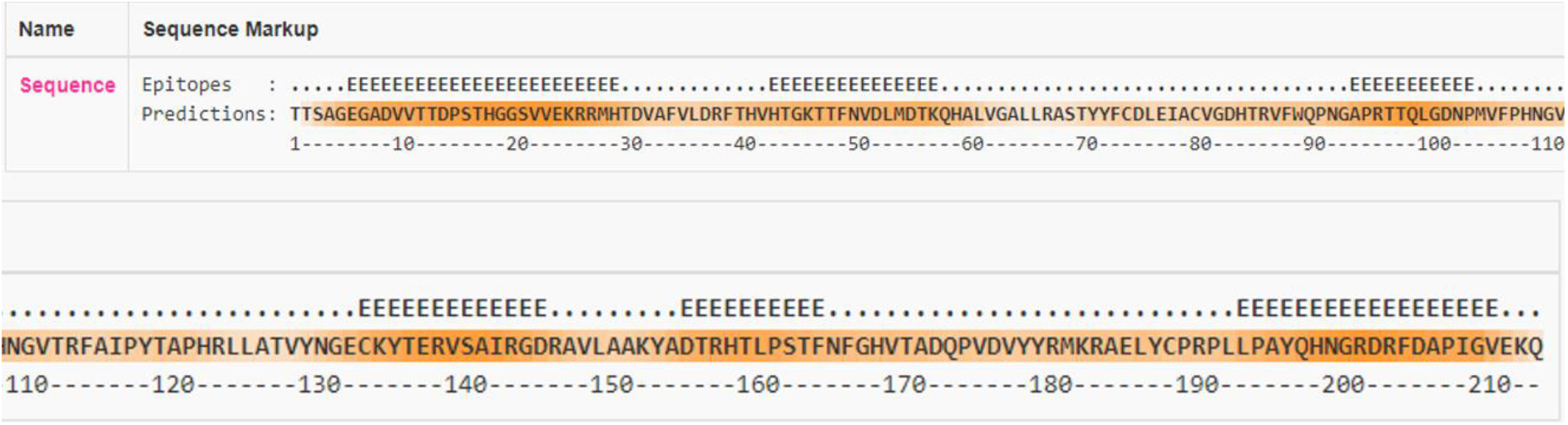
Representation of sequential B-cell epitopes prediction

The 5ACA VP1 based model was selected for a 3D presentation and the consensus sequence aligned to the 5ACA VP1 sequence to study the variability results are presented in Table 1 with the predicted B-cell epitopes.

Furthermore, discontinuous B-cell epitopes were determined using the IEDB online platform by setting the threshold at 0.5. The resulting eight peptides are represented in Table 2:

**Table 2:** Predicted Discontinuous Epitopes

| No | Residues | Number of<br>residues | Score |
| --- | --- | --- | --- |
| 1 | T136, T137, Q138 | 3 | 0.929 |
| 2 | T1, T2, S3, S4, G5, E6, G7, A8, D9, V10, V11, T12, T13, D14, P15, S16, T17, H18, G19, G20, A21, V22, T23, E24 | 24 | 0.854 |
| 3 | P198, G199, Y200, D201, H202, A203, D204, R205, D206, R207, F208, D209, S210, P211, I212, G213 | 16 | 0.816 |
| 4 | Y130, N131, G132, E133, C134, K135 | 6 | 0.806 |
| 5 | T44, N45, T47, L81, G82, E83, H84, E85, R86, W88, T97, T98, T99, L100, R101, D102, N103, P104, M105, V106, F107, S108, H109, N110, N111, V112, A175, D176, K177, P178 | 30 | 0.645 |
| 6 | P91, N92, G93, A94, P95, R96 | 6 | 0.562 |

**Table 3:**
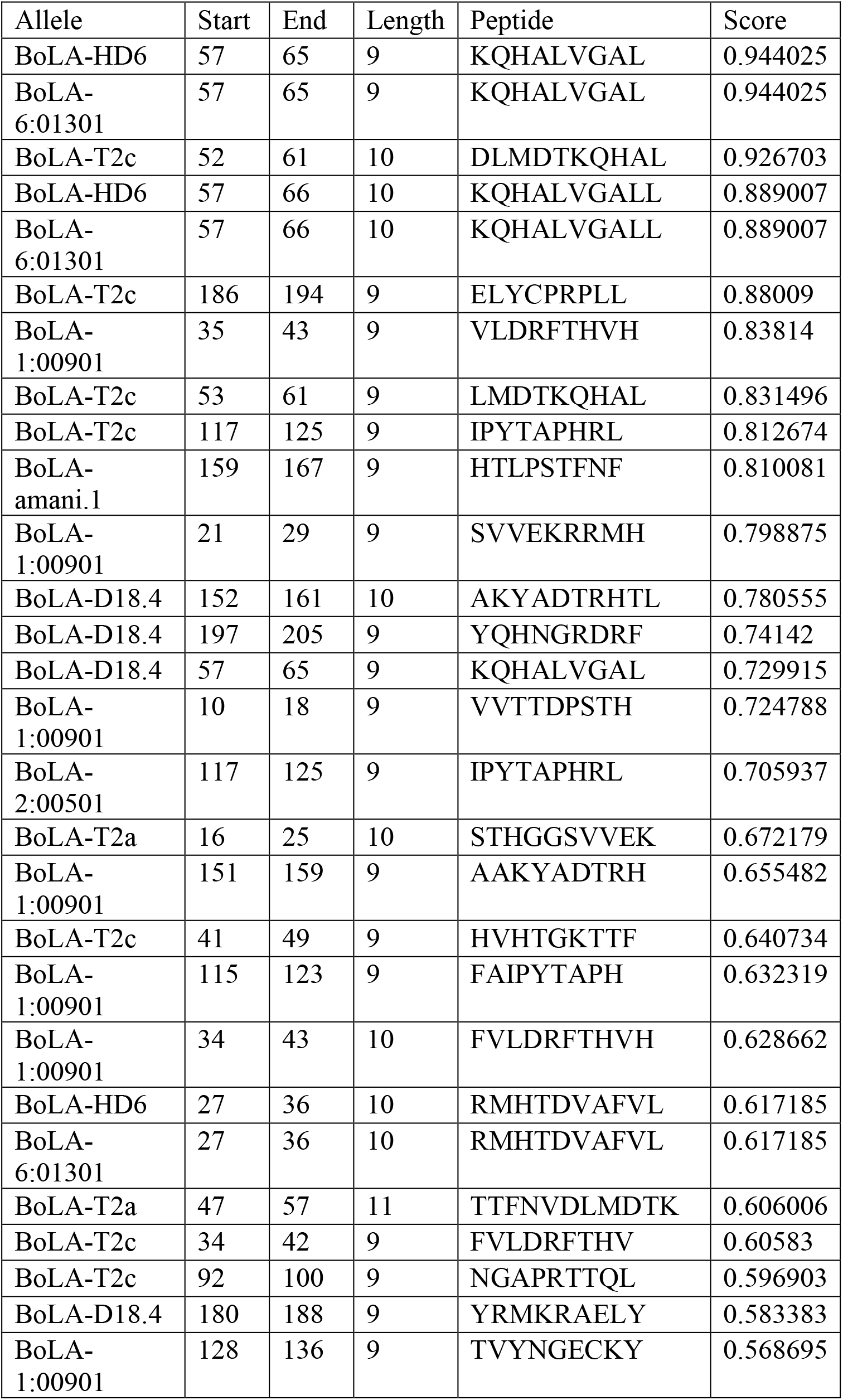
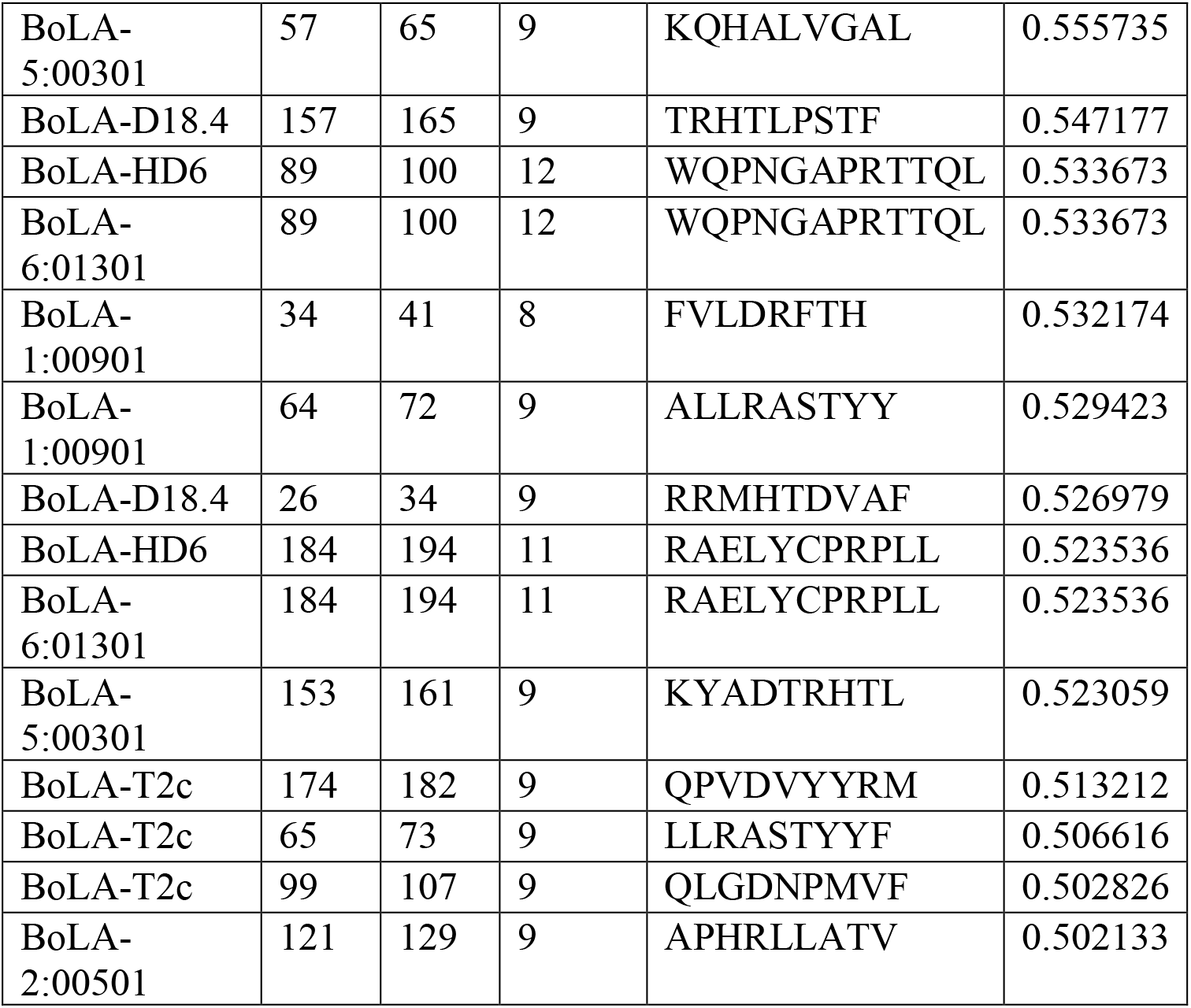
Different combinations of BoLA alleles with several peptides with their scores

Using the PyMol software, we loaded the model (PDB ID:5aca) and mapped both linear and discontinuous epitopes. For visibility, we only present epitopes scoring ≥ 8.

#### 3.2.2. T-cell epitopes

The following alleles were selected (BoLA-1:00901, BoLA-2:00501, BoLA-3:00101, BoLA-4:02401, BoLA-5:00301, BoLA-6:01301, BoLA-amani.1, BoLA-AW10, BoLA-D18.4, BoLA-gb1.7, BoLA-HD6, BoLA-JSP.1, BoLA-T2a, BoLA-T2b, BoLA-T2c, BoLA-T5, BoLA-T7) to analyze their affinity to different peptide sizes ranging from 8 to 12 amino acids. By considering the threshold of score ≥ 0.5, we remained with 42 potential epitopes (Table 2). Among the latter, at least ten pairs (BoLA-HD6*KQHALVGAL, BoLA-6:01301*KQHALVGAL, BoLA-T2c*DLMDTKQHAL, BoLA-HD6*KQHALVGALL, BoLA-6:01301*KQHALVGALL, BoLA-T2c*LMDTKQHAL, BoLA-T2c*IPYTAPHRL and BoLA-amani.1*HTLPSTFNF) scored more than 0.8. Furthermore, certain sequence regions contained peptides that could be considered to induce both B-cell and T-cell immune responses, among them two pairs (BoLA-1:00901*SVVEKRRMH and BoLA-amani.1*HTLPSTFNF) had scored 0.798 and 0.810 consecutively. The epitope in the second couple (res 156-166) is shown to be very variable which can lead to vaccine failure and is located in a region able to induce both B-cell and T-cell immunity.

Furthermore, we joined several epitopes by removing the noise and we adopted the GPGPG spacer as previously used (17). However, some epitopes overlapped and were combined as continuous residue. Therefore, the resulting multiepitope peptide is ^10^VVTTDPSTHGGSVVEKRRMHTDVAFVLDRFTHVHTGKTTFNVDLMDTKQHALVG ALLRASTYYF^73^GPGPG^89^WQPNGAPRTTQLGDNPMVF^107^GPGPG^115^FAIPYTAPHRLLA TVYNGECKY^136^*GPGPG*^151^AAKYADTRHTLPSTFNF^167^. Finally, we mapped epitopes scoring ≥8 to the 3D model (figure 7).

**Figure 6:**
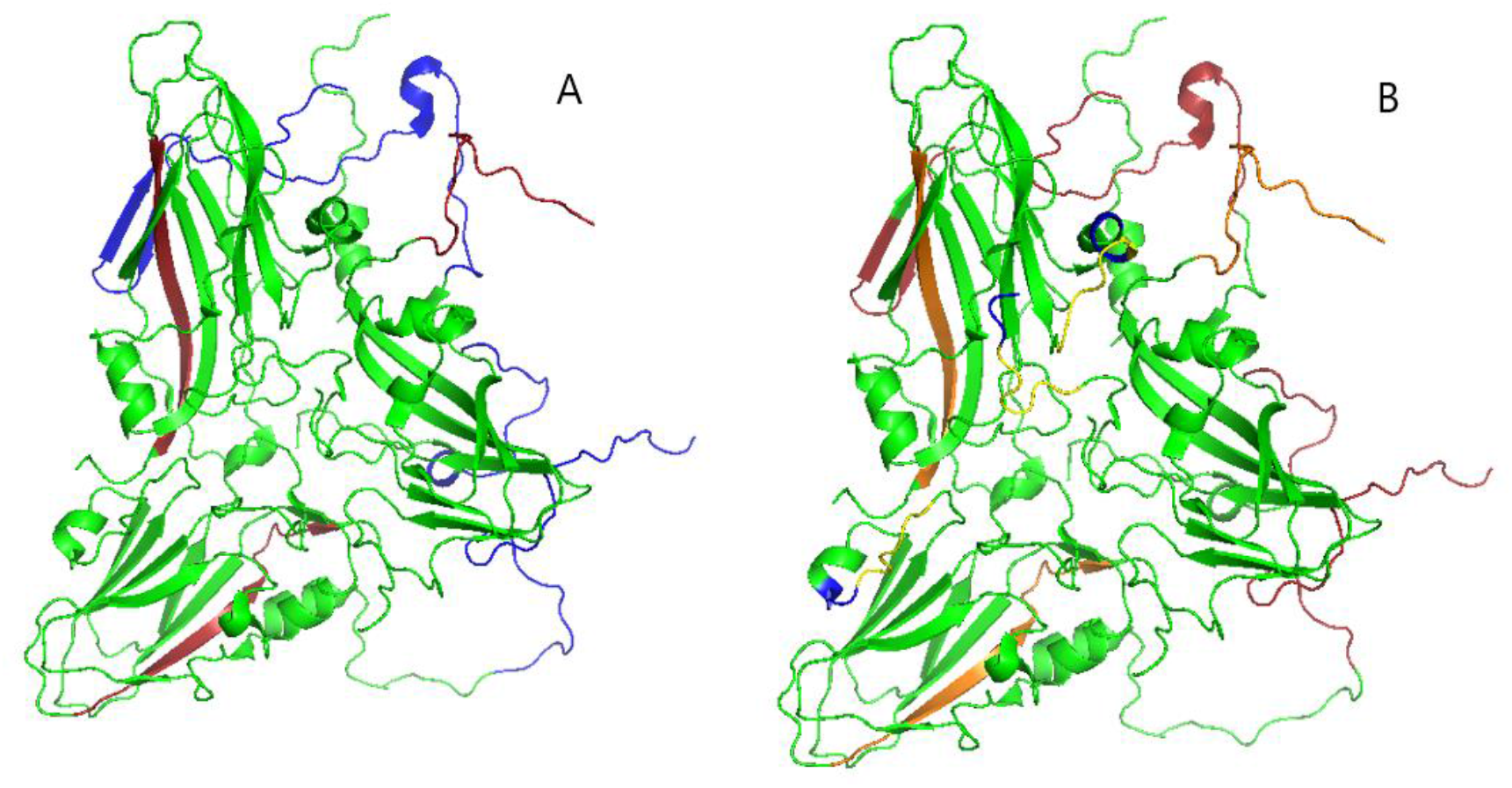
Humoral epitope prediction on the FMDV SAT2 VP1 consensus sequence isolated in East Africa. Fig. 6.A represents linear epitopes with residue 198-213 (firebrick) and residue 1-30 (blue) while Fig6.B represents discontinuous epitopes with residue 136-138 (blue), residue 1-24 (firebrick), residue 198-213 (orange) and residue 130-135 (yellow).

**Figure 7:**
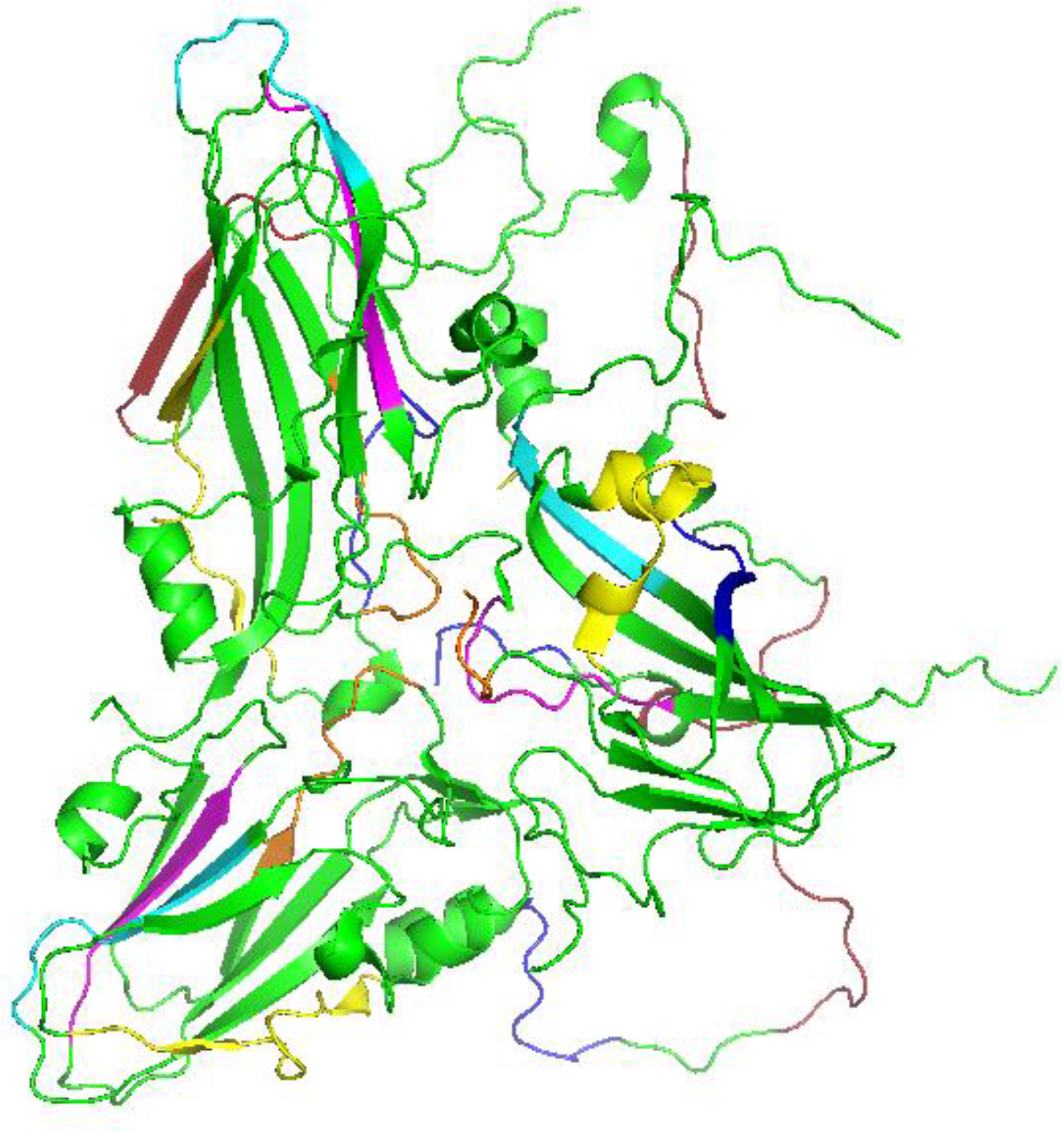
Three-dimensional mapping on the selected model (PDB ID: 5aca) of the predicted MHC I T-cell epitopes scoring ≥8. Epitope residues are highlighted as follows: residue 21-29 (firebrick), residue 35-43 (blue), residue 52-66 (yellow), residue 117-125 (magenta), residue 159-167 (orange) and residue 186-194 (cyan).

## 4. Discussion

The present study proposes peptides with a higher probability to induce B and T cell immune responses in cattle and therefore can be considered as good candidates for vaccine design. Using the Wu-Kabat variability index available at the protein variability server, the variability plot we obtained with SAT2 was less variable compared to the Asia1 serotype variability obtained by Sultana *et al.*(18). This study emphasizes a regional intratypic variability found at least in three positions. Sahle *et al.*(19) described variable sites (positions 46-51, 109-114, 134-147 and 152-157) in SAT 1 that are approximately equivalent to the ones observed with this study’s SAT 2 sequences (positions 45-50, 107-111, 135-141 and 151-160) with the RGD site (149-151) and C- terminus (202-217) being relatively conserved for both serotypes.

Though *in silico* prediction of epitopes is very important, it has to be coupled with thermo-stabilization of the identified epitopes because of the thermal lability of SAT serotypes (20). The BoLA system showed to have an affinity to surface-exposed peptides of the VP1 with the BoLA-HD6 allele to have the highest affinity while Agrawal *et al.* found that BoLA-T7 had the highest percentile rank towards PPRV peptides (21). The identified epitopes can be used to design a polytope-based plasmid DNA vaccine using linkers and adjuvant (17,22) that can be tested in *vitro* and *in vivo*.

## 5. Conclusion

Experimentations using the predicted peptides can lead to region-tailored DNA vaccines development. Special focus should be put on residues that showed low variability and to induce B-cell immunity and high affinity to BoLA alleles for *in vitro* and *in vivo* testing. A deeper understanding of molecular epidemiology in East Africa can improve the effectiveness of the predicted epitopes. We, therefore, recommend more fieldwork especially in countries where little is known in terms of circulating strains. In the future, advanced AI-based tools to predict protein folding, such as Alpha Fold 2, can be used for epitope mapping on 3D structures.

## Supporting information

Supplemental Sequences

## References

1. Stanway G, Brown F, Christian PD, Hovi T, Hyypiä T, King AMQ, et al. Taxonomy of the Picornaviridae?: Species Designations and Three New Genera. Abstracts of the XIth Meeting of the European Study Group on Molecular Biology of Picornaviruses. [Internet]. 2000. Available from: https://www.picornastudygroup.com/posters/europic_2000.pdf

2. Jamal SM, Belsham GJ. Foot-and-mouth disease: past, present and future. Vet Res. 2013;44(116):1–14.

3. Sumption K, Rweyemamu M, Wint W. Incidence and Distribution of Foot-and-Mouth Disease in Asia, Africa and South America; Combining Expert Opinion, Official Disease Information and Livestock Populations to Assist Risk Assessment. Transbound Emerg Dis [Internet]. 2008 Jun 28;55(1):5–13. Available from: https://onlinelibrary.wiley.com/doi/10.1111/j.1865-1682.2007.01017.x

4. Brito BP, Rodriguez LL, Hammond JM, Pinto J, Perez AM. Review of the Global Distribution of Foot-and-Mouth Disease Virus from 2007 to 2014. Transbound Emerg Dis [Internet]. 2017 Apr;64(2):316–32. Available from: https://onlinelibrary.wiley.com/doi/10.1111/tbed.12373

5. Udahemuka JC, Aboge GO, Obiero GO, Lebea PJ, Onono JO, Paone M. Risk factors for the incursion, spread and persistence of the foot and mouth disease virus in Eastern Rwanda. BMC Vet Res. 2020 Dec;16(1):387.

6. Rowlands DJ, Clarke BE, Carroll AR, Brown F, Nicholson BH, Bittle JL, et al. Chemical basis of antigenic variation in foot-and-mouth disease virus. Nature [Internet]. 1983 Dec;306(5944):694–7. Available from: http://www.nature.com/articles/306694a0

7. Waters R, Ludi AB, Fowler VL, Wilsden G, Browning C, Gubbins S, et al. Efficacy of a high-potency multivalent foot-and-mouth disease virus vaccine in cattle against heterologous challenge with a field virus from the emerging A/ASIA/G-VII lineage. Vol. 36, Vaccine. 2018. p. 1901–7.

8. FAO/EuFMD. EuFMD P II: European Neighborhood/ Report on Significant FAST disease events and information [Internet]. Rome; 2020. Available from: https://rr-middleeast.oie.int/wp-content/uploads/2021/01/eufmd_pii_report_on_significant_fast_disease_events_and_information_october_december_2020.pdf

9. Wang CY, Chang TY, Walfield AM, Ye J, Shen M, Chen SP, et al. Effective synthetic peptide vaccine for foot-and-mouth disease in swine. Vaccine [Internet]. 2002 Jun;20(19–20):2603–10 Available from: https://linkinghub.elsevier.com/retrieve/pii/S0264410X02001482

10. Forner M, Cañas-Arranz R, Defaus S, de León P, Rodríguez-Pulido M, Ganges L, et al. Peptide-Based Vaccines: Foot-and-Mouth Disease Virus, a Paradigm in Animal Health. Vaccines [Internet]. 2021 May 8;9(5):477. Available from: https://www.mdpi.com/2076-393X/9/5/477

11. Defaus S, Forner M, Cañas-Arranz R, de León P, Bustos MJ, Rodríguez-Pulido M, et al. Designing Functionally Versatile, Highly Immunogenic Peptide-Based Multiepitopic Vaccines against Foot-and-Mouth Disease Virus. Vaccines [Internet]. 2020 Jul 22;8(3):406. Available from: https://www.mdpi.com/2076-393X/8/3/406

12. Kabat EA, Wu TT, Bilofsky H. Unusual distributions of amino acids in complementarity determining (hypervariable) segments of heavy and light chains of immunoglobulins and their possible roles in specificity of antibody-combining sites. J Biol Chem. 1977;252(19):6609–16.

13. Garcia-Boronat M, Diez-Rivero CM, Reinherz EL, Reche PA. PVS: a web server for protein sequence variability analysis tuned to facilitate conserved epitope discovery. Nucleic Acids Res [Internet]. 2008 May 19;36(Web Server):W35–41. Available from: https://academic.oup.com/nar/article-lookup/doi/10.1093/nar/gkn211

14. Reynisson B, Alvarez B, Paul S, Peters B, Nielsen M. NetMHCpan-4.1 and NetMHCIIpan-4.0: improved predictions of MHC antigen presentation by concurrent motif deconvolution and integration of MS MHC eluted ligand data. Nucleic Acids Res [Internet]. 2020 Jul 2;48(W1):W449–54. Available from: https://academic.oup.com/nar/article/48/W1/W449/5837056

15. Schrödinger L. The PyMOL Molecular Graphics System, Version 2.5. 2021.

16. Galgonek J, Vymětal J, Jakubec D, Vondrášek J. Amino Acid Interaction (INTAA) web server. Nucleic Acids Res [Internet]. 2017 Jul 3 [cited 2021 Aug 6];45(W1):W388–92. Available from: http://www.cbs.dtu.dk/services/BepiPred/

17. Raza S, Siddique K, Rabbani M, Yaqub T, Anjum AA, Ibrahim M, et al. In silico analysis of four structural proteins of aphthovirus serotypes revealed significant B and T cell epitopes. Microb Pathog [Internet]. 2019;128(January):254–62. Available from: https://doi.org/10.1016/j.micpath.2019.01.007

18. Sultana M, Alam S, Rahmman MZ, Hossain M, Amin R. Antigenic heterogeneity of capsid protein VP1 in foot-and-mouth disease virus (FMDV) serotype Asia1. Adv Appl Bioinforma Chem [Internet]. 2013 Aug;6(1):37. Available from: http://www.dovepress.com/antigenic-heterogeneity-of-capsid-protein-vp1-in-foot-and-mouth-diseas-peer-reviewed-article-AABC

19. Sahle M, Dwarka RM, Venter EH, Vosloo W. Comparison of SAT-1 foot-and-mouth disease virus isolates obtained from East Africa between 1971 and 2000 with viruses from the rest of sub-Saharan Africa. Arch Virol [Internet]. 2007 Apr 3;152(4):797–804. Available from: http://link.springer.com/10.1007/s00705-006-0893-x

20. Doel TR, Baccarini PJ. Thermal stability of foot-and-mouth disease virus. Arch Virol [Internet]. 1981 Mar;70(1):21–32. Available from: http://link.springer.com/10.1007/BF01320790

21. Agrawal A, Gupta R, Gattani A, Patel SK, Khan MH, Singh P. Novel T Cell Epitope Designing from PPRV HN Protein for Peptide based Subunit Vaccine: An Immune Informatics Approach. Int J Curr Microbiol Appl Sci [Internet]. 2020 Mar 20;9(3):2432–9. Available from: https://www.ijcmas.com/abstractview.php?ID=16563&vol=9-3-2020&SNo=278

22. Michel-Todó L, Bigey P, Reche PA, Pinazo M-J, Gascón J, Alonso-Padilla J. Design of an Epitope-Based Vaccine Ensemble for Animal Trypanosomiasis by Computational Methods. Vaccines [Internet]. 2020 Mar 16;8(1):130. Available from: https://www.mdpi.com/2076-393X/8/1/130

